# Melanin-concentrating hormone signaling regulates prefrontal updating of learned avoidance

**DOI:** 10.64898/2026.06.26.734738

**Authors:** Pratheba Kandasamey, Eva Bracey, Ladina Odermatt, Denis Burdakov, Daria Peleg-Raibstein

**Affiliations:** ETH Zürich: Department of Health Sciences and Technology (D-HEST); Institute for Neuroscience; Neuroscience Center Zürich (ZNZ)

**Keywords:** active avoidance, extinction, melanin-concentrating hormone, medial prefrontal cortex, instrumental defensive behavior

## Abstract

Adaptive avoidance depends on a delicate balance: animals must act rapidly when a cue predicts danger, but suppress the same action when the cue no longer has consequence. How MCH neuromodulatory signaling shapes this prefrontal updating process remains poorly understood. Here, we identify melanin-concentrating hormone receptor 1 (MCHR1) signaling as a regulator of active avoidance extinction. Pharmacological MCHR1 antagonism with SNAP-94847 left acquisition of two-way active avoidance intact, but promoted extinction once the tone was no longer followed by shock. This effect was reproduced by prelimbic mPFC-targeted SNAP infusion, indicating that prefrontal MCHR1 signaling contributes to the persistence of learned avoidance. Fiber photometry from CaMKII-positive mPFC neurons revealed that MCHR1 antagonism enhanced excitatory prefrontal activity during successful avoidance and altered trial-history-dependent mPFC activity during extinction, most prominently on avoidance trials that followed previous avoidance. These findings identify MCHR1 signaling as a regulator of adaptive avoidance updating and suggest that MCHR1 antagonism facilitates extinction by altering prefrontal processing of recent action history when a formerly protective response loses behavioral value.

**Significance Statement:** In anxiety- and trauma-related disorders, avoidance can persist long after danger is gone, interfering with daily life. While avoidance is essential for survival, it can become harmful when it is no longer needed. We found that blocking brain receptors for melanin-concentrating hormone helps mice stop responding to outdated warning signals while preserving their ability to learn from danger. We identify the medial prefrontal cortex as a key brain region where this intervention changes activity during fear-guided behavior. This study highlights a potential therapeutic strategy for reducing excessive avoidance without compromising normal protective responses.

## Introduction

Adaptive defensive behavior requires animals not only to learn which cues predict threat, but also to update defensive actions when those cues lose their consequence. Active avoidance provides a powerful framework for studying this process because it combines Pavlovian threat prediction with instrumental action control: animals use a warning cue to perform an action that prevents an aversive outcome, and during extinction they must suppress that action when it is no longer required (1–5) (6). Whereas Pavlovian freezing reflects a passive defensive reaction to predicted threat, avoidance requires the animal to assign behavioral significance to an action that changes the expected outcome and then update this cue–action–outcome relationship when the contingency changes. Persistent avoidance and safety behaviors are clinically important because they can maintain anxiety and trauma-related symptoms by preventing corrective learning that previously threatening cues can become safe (7–14).

The medial prefrontal cortex (mPFC) is central to this form of behavioral control. More broadly, medial prefrontal and frontostriatal circuits support behavioral flexibility and updating of action strategies when contingencies change (15–19). Prefrontal circuits regulate the balance between reactive defensive responses, instrumental coping strategies and extinction when threat contingencies change (6);(20). Work in active avoidance has shown that prelimbic and infralimbic mPFC subregions, together with striatal and amygdala circuits, make dissociable contributions to avoidance expression, action selection and extinction (15, 20–24); (25); (26); (27). This is particularly relevant for the present study because recent population-imaging work in the prelimbic mPFC showed that excitatory prefrontal activity encodes avoidance-specific action patterns that predict tone-induced avoidance and are separable from spontaneous movements with similar kinematics (26, 28).

Despite this progress, the neuromodulatory systems that tune mPFC-dependent avoidance updating remain poorly understood. Melanin-concentrating hormone (MCH) neurons, located mainly in the lateral hypothalamus and zona incerta, have classically been studied in relation to energy balance, feeding, sleep, and reward (29–34). However, accumulating evidence indicates that the MCH system also regulates memory and emotional flexibility (35–37). Hypothalamic MCH neurons are required for normal fear extinction and long-term flexibility of fear behavior (38), while REM-active MCH neurons promote forgetting of hippocampus-dependent memories (39). MCH signaling has also been implicated in anxiety-like behavior and impulsive action control, suggesting a broader role in adapting behavior to changing internal and external conditions (39);(40). Thus, the field has established that MCH signaling can influence affective state, memory persistence and behavioral flexibility, but it remains unknown whether its receptor MCHR1 regulates active avoidance extinction, where animals must update a learned cue–action–outcome relationship through prefrontal control mechanisms.

Together, these observations define a specific gap: MCH signaling has been linked to emotional flexibility and memory persistence, and mPFC circuits are known to control avoidance updating, but it remains unknown whether MCHR1 signaling links these systems during extinction of learned avoidance. This question is particularly relevant because extinction depends on using recent non-reinforced cue exposures to revise whether an avoidance action remains necessary, a process strongly associated with mPFC function. We also examined aged animals as a boundary condition, because aging compromises cognitive flexibility and alters mPFC excitability, raising the possibility that MCHR1-dependent avoidance updating may be reduced when prefrontal flexibility is limited (41); (42); (43); (44).

Here, we tested whether MCHR1 antagonism regulates acquisition and extinction of two-way active avoidance and whether this regulation involves excitatory activity in the prelimbic region of the mPFC. We combined systemic SNAP-94847 administration, aged-mouse behavior, prelimbic mPFC-targeted SNAP infusion, and CaMKII-driven mPFC fiber photometry during avoidance acquisition, extinction, and trial-history-defined behavioral transitions. By recording CaMKII-positive mPFC neurons, we focused on excitatory pyramidal populations that form the principal cortical output through which mPFC can influence downstream avoidance circuits. This approach allowed us to test whether MCHR1 signaling contributes to the neural control of adaptive defensive action and recent-outcome updating.

## Results

### Systemic MCHR1 antagonism facilitates extinction of active avoidance in adult mice

We first tested whether systemic MCHR1 antagonism alters acquisition or extinction of two-way active avoidance. Adult mice received SNAP-94847 or vehicle during a shuttle-box task in which crossing to the opposite compartment during the tone prevented footshock during acquisition; during extinction, the tone was presented without shock (Fig. 1a-c). SNAP- and vehicle-treated mice acquired avoidance across training and reached comparable avoidance levels by the end of acquisition (Fig. 1d; two-way repeated-measures ANOVA: 10-trial bins F(15,210) = 29.46, p < 0.0001; Group and 10-trial bins x Group interaction, n.s.), indicating that systemic MCHR1 antagonism did not impair learning of the tone-shock contingency or execution of the shuttle response. During extinction, avoidance decreased across days in both groups, but SNAP-treated mice showed significantly lower avoidance than controls (Fig. 1e; 10-trial bins, F(11,154) = 8.758, p < 0.0001; Group, F(1,14) = 13.72, p = 0.002; 10-trial bins × Group, F(11,154) = 0.9204, p = 0.5225). This indicates that systemic MCHR1 antagonism facilitated extinction of the instrumental avoidance response without affecting acquisition.

**Fig. 1.**
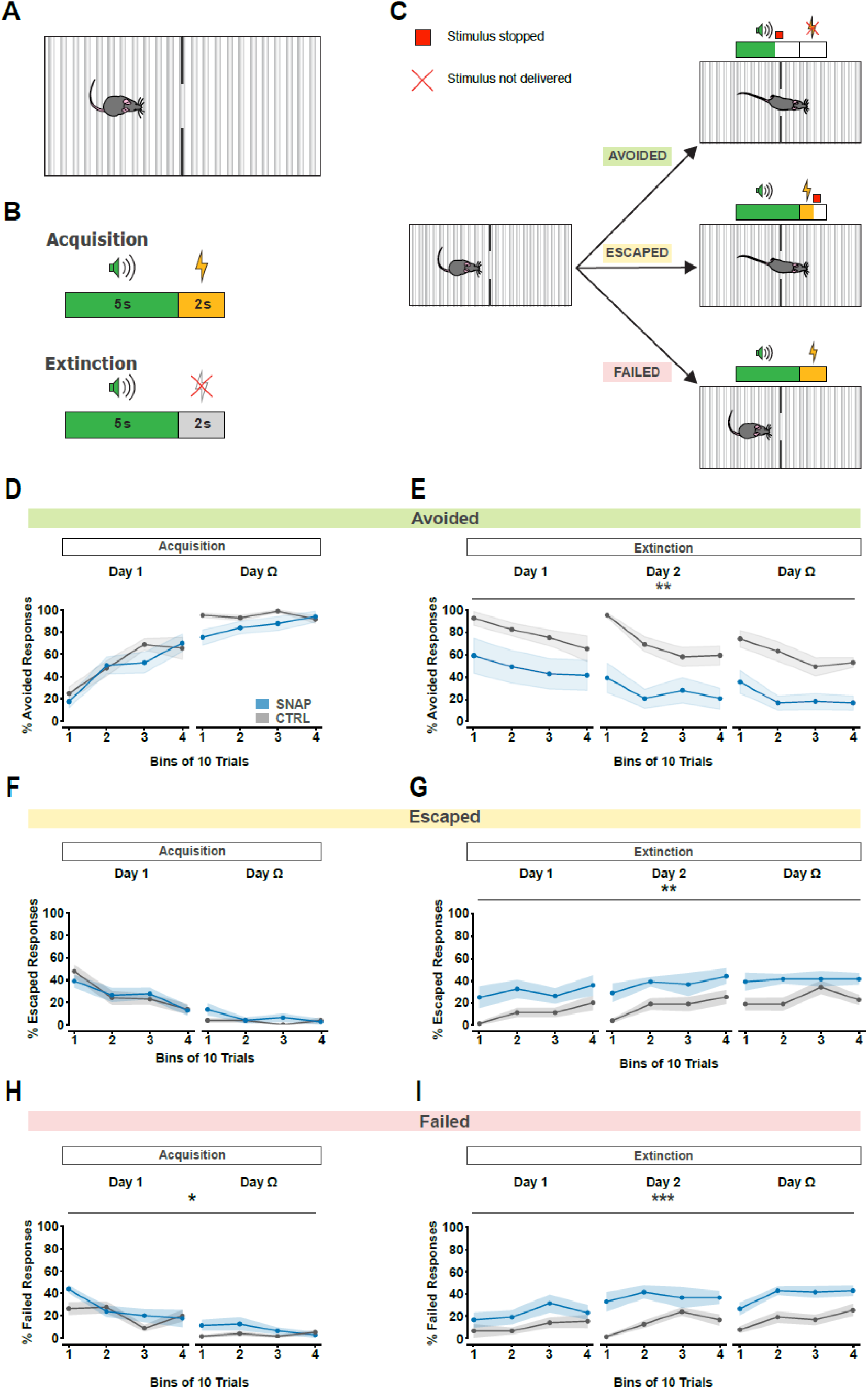
Systemic MCHR1 antagonism facilitates extinction of active avoidance in adult mice. **a.** Schematic top-view representation of the experimental apparatus used for two-way active avoidance testing. The chamber consisted of two compartments equipped with shock grids and separated by a central wall containing an opening that allowed the mouse to move freely between compartments. A speaker was positioned within the apparatus to deliver the conditioned stimulus (CS; cue tone). The maximum CS (tone) duration is 5 s and the maximum US (shock) duration is 2 s. **b.** Task schematic representing acquisition and extinction. In both phases the CS was presented; during acquisition it was followed by shock, whereas during extinction the shock was omitted. **c.** Trial structure of acquisition trials and illustration of the different trial outcomes (avoided, escaped, failed). A compartment change during the CS terminated the tone and was scored as avoided. A compartment change during the US terminated the shock and was scored as escaped. No compartment change during the CS or US was scored as failed. Extinction trials were identical except that the shock was omitted. **d.** Avoided responses across acquisition. Two-way repeated-measures ANOVA revealed a significant main effect of 10-trial bins (F(15, 210) = 29.46, p < 0.0001), with no significant Group effect (F(1,14) = 1.99, p = 0.18 or 10-trial bins × Group interaction (F(15,210) = 1.43, p = 0.14. **e.** % Avoided responses across extinction. Two-way repeated-measures ANOVA revealed significant main effects of 10-trial bins (F(11,154) = 8.78, p < 0.0001) and Group factor (F(1,14) = 13.72, p = 0.002), with no significant interaction between factors (F(11,154) = 0.92, p = 0.52). **f.** % Escaped responses across the days within acquisition phase. Two-way repeated-measures ANOVA revealed a significant main effect of 10-trial bins (F(15,210) = 18.69, p < 0.0001), with no significant main effect of Group factor (F(1,14) = 0.24, p = 0.63) and no significant interaction between factors (F(15, 210) = 1.05, p = 0.41). **g.** % Escaped responses across extinction. Two-way repeated-measures ANOVA revealed significant main effects of 10-trial bins factor (F(11,154) = 4.44, p = 0.001) and Group factor (F(1,14) = 9.23, p = 0.009), with no significant interaction between factors (F(11,154) = 0.46, p = 0.93). **h.** % Failed responses across acquisition. Two-way repeated-measures ANOVA revealed significant main effects of 10-trial bins factor (F(15,210) = 12.21, p < 0.0001) and Group factor (F(1,14) = 5.31, p = 0.04), with no significant interaction between factors (F(15,210) = 1.58, p = 0.08). **i.** % Failed responses across the days within extinction phase. Two-way repeated-measures ANOVA revealed significant main effects of 10-trial bins factor (F(11,154) = 4.57, p = 0.001) and Group factor (F(1,14) = 19.66, p = 0.0006), with no significant interaction between factors (F(11,154) = 1.09, p = 0.37). Data are shown as mean ± s.e.m.; n = 8 mice in control group (5 males and 3 females), n = 8 mice in SNAP group (5 males and 3 females)

Because each trial could end in one of three mutually exclusive outcomes, escaped and failed responses provide complementary information about how changes in avoidance were expressed: during acquisition, escapes reflect delayed but successful responses after shock onset, whereas failures reflect absence of a shuttle response; during extinction, when shock is omitted, failures instead indicate that the animal withheld the previously reinforced avoidance action. During acquisition, escaped responses declined across training without a group effect (Fig. 1f; 10-trial bins, F(15,210) = 18.69, p < 0.0001; Group, F(1,14) = 0.24, p = 0.63; 10-trial bins × Group, F(15,210) = 1.05, p = 0.40), supporting intact acquisition of avoidance responding under SNAP. During extinction, escaped responses differed between groups (Fig. 1g; Group, F(1,14) = 9.23, p = 0.009), and failed responses were also increased under SNAP (Fig. 1i; Group, F(1,14) = 19.66, p = 0.0006). Thus, the reduced avoidance in SNAP-treated mice reflected a redistribution away from cue-driven shuttle responses after shock omission, rather than impaired acquisition or inability to perform the action. Together, these data show that systemic MCHR1 antagonism facilitates extinction by reducing persistent cue-driven avoidance after shock omission.

### Systemic MCHR1 antagonism transiently facilitates early extinction in aged mice

We next tested whether the extinction-promoting effect of systemic MCHR1 antagonism generalizes to aged mice. This experiment was motivated by evidence that aging reduces extinction and cognitive flexibility and alters mPFC excitability, raising the possibility that MCHR1-dependent modulation of avoidance updating may be weakened in aged animals. Aged mice received SNAP-94847 or vehicle and were trained in the same two-way active avoidance framework used for the adult cohort, with acquisition followed by extinction. Both groups acquired the avoidance response across training, with a significant effect of 10-trial bins but no group effect or 10-trial bins × group interaction (Fig. 2a; two-way repeated-measures ANOVA: 10-trial bins, F(7,189) = 49.53, p < 0.0001; Group, F(1,27) = 0.35, p = 0.56; 10-trial bins × Group, F(7,189) = 0.73, p = 0.64). Thus, systemic MCHR1 antagonism did not impair acquisition of the instrumental avoidance response in aged mice. Across the full extinction phase, avoidance decreased over days but did not show a sustained group difference (Fig. 2b; 10-trial bins, F(7,189) = 12.20, p < 0.0001; Group, F(1,27) = 0.81, p = 0.38; 10-trial bins × Group, F(7,189) = 1.31, p = 0.25). However, a focused analysis of the first extinction day revealed a significant group difference (Welch’s t-test: t(5.93) = 6.24, p = 0.0008).

**Fig. 2.**
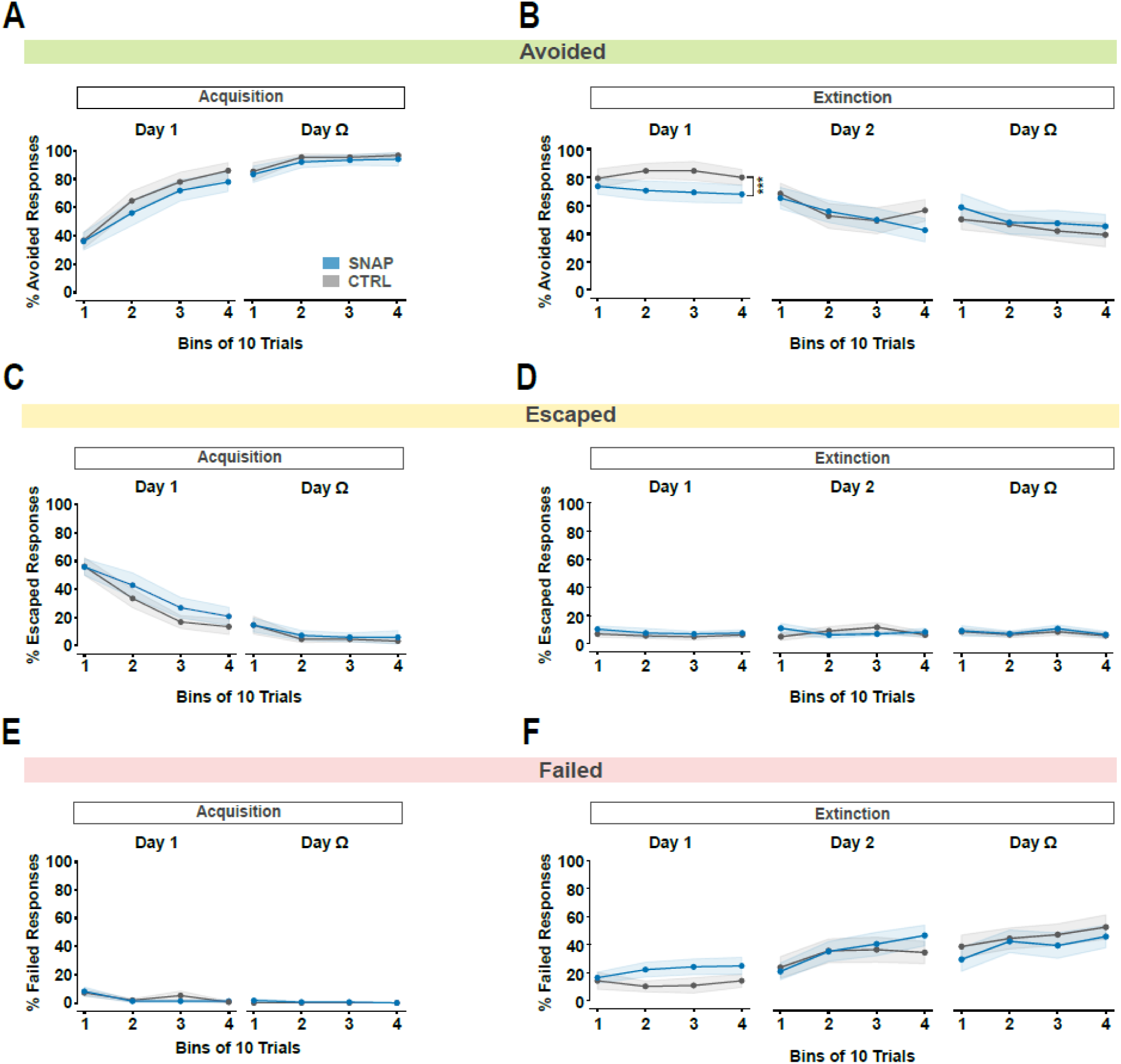
Systemic MCHR1 antagonism transiently facilitates early extinction in aged mice. **a.** % Avoided responses across the days within acquisition phase. Two-way repeated-measures ANOVA revealed a significant main effect of 10-trial bins (F(7,189) = 49.53, p < 0.0001), with no significant main effect of Group factor (F(1,27) = 0.35, p = 0.56) and no significant interaction between factors (F(7,189) = 0.73, p = 0.64)**. b** % Avoided responses across days within extinction phase. Two-way repeated-measures ANOVA revealed a significant main effect of 10-trial bins (F(7,189) = 12.20, p < 0.0001), with no significant main effect of Group factor (F(1,27) = 0.81, p = 0.38) and no significant interaction between factors (F(7,189) = 1.31, p = 0.25). Zoomed-in analysis of the first extinction day using an unpaired two-tailed Welch’s t-test revealed a significant difference between SNAP and control groups (t(5.934) = 6.24, p = 0.0008). **c.** % Escaped responses across the days within acquisition phase. Two-way repeated-measures ANOVA revealed a significant main effect of 10-trial bins (F(7,189) = 33.09, p < 0.0001), with no significant main effect of Group factor (F(1,27) = 0.43, p = 0.52) and no significant interaction between factors (F(7, 189) = 2.04, p = 0.05). **d**. % Escaped responses across the days within extinction phase. Two-way repeated-measures ANOVA revealed no significant main effects of 10-trial bins (F(11,297) = 0.95, p = 0.49) or Group factor (F(1,27) = 0.43, p = 0.52), and no significant interaction between factors (F(11,297) = 0.73, p = 0.71). **e.** % Failed responses across the days within acquisition phase. Two-way repeated-measures ANOVA revealed a significant main effect of 10-trial bins (F(7,189) = 5.86, p < 0.0001), with no significant main effect of Group factor (F(1,27) = 0.06, p = 0.81) and no significant interaction between factors (F(7,189) = 1.01, p = 0.42). **f**. % Failed responses across the days within extinction phase. Two-way repeated-measures ANOVA revealed a significant main effect of 10-trial bins (F(11,297) = 11.28, p < 0.0001), with no significant main effect of Group factor (F(1,27) = 0.04, p = 0.83) and no significant interaction between factors (F(11, 297) = 1.44, p = 0.16). Data are shown as mean ± s.e.m.; n = 15 mice in control group (8 males and 7 females), n = 14 mice in SNAP group (7 males and 7 females)

We next examined escaped and failed responses to determine whether the early reduction in avoidance was accompanied by a broader redistribution of behavioral outcomes. Escaped responses changed during acquisition but did not differ between groups during acquisition or extinction (Fig. 2c,d; acquisition Group, F(1,27) = 0.43, p = 0.52; extinction Group, F(1,27) = 0.43, p = 0.52). Failed responses also changed across days but were not significantly affected by SNAP during either acquisition or extinction (Fig. 2e,f; acquisition Group, F(1,27) = 0.06, p = 0.81; extinction Group, F(1,27) = 0.04, p = 0.83). These data indicate that, unlike in adult mice, systemic MCHR1 antagonism produced only a temporally restricted early reduction in avoidance in aged animals, without a sustained extinction-related redistribution across behavioral outcomes. Thus, aging may limit the persistence of MCHR1-dependent modulation of avoidance extinction.

### mPFC-targeted MCHR1 antagonism facilitates extinction of active avoidance

We next asked whether MCHR1 signaling in the medial prefrontal region contributes to active avoidance extinction. This experiment was designed to test whether local prefrontal MCHR1 antagonism is sufficient to modulate avoidance when threat contingencies change. Mice were implanted with bilateral cannulae targeting the mPFC and received local infusion of SNAP-94847 or vehicle before testing in the two-way active avoidance task (Fig. 3a-c). During acquisition, mPFC-targeted SNAP did not reduce avoidance relative to vehicle controls (Fig. 3d; 10-trial bins, F(9,72) = 5.16, p < 0.0001; Group, F(1,8) = 0.038, p = 0.85; 10-trial bins × Group, F(9,72) = 1.80, p = 0.08). Thus, local MCHR1 antagonism did not impair acquisition or performance of the avoidance response. During extinction, mPFC-targeted SNAP significantly reduced avoidance (Fig. 3e; Group, F(1,8) = 9.55, p = 0.015; 10-trial bins, F(29,232) = 2.19, p = 0.0008; 10-trial bins × Group, F(29,232) = 1.25, p = 0.19). This finding supports a role for MCHR1 signaling in the medial prefrontal region in regulating extinction of active avoidance.

**Fig. 3.**
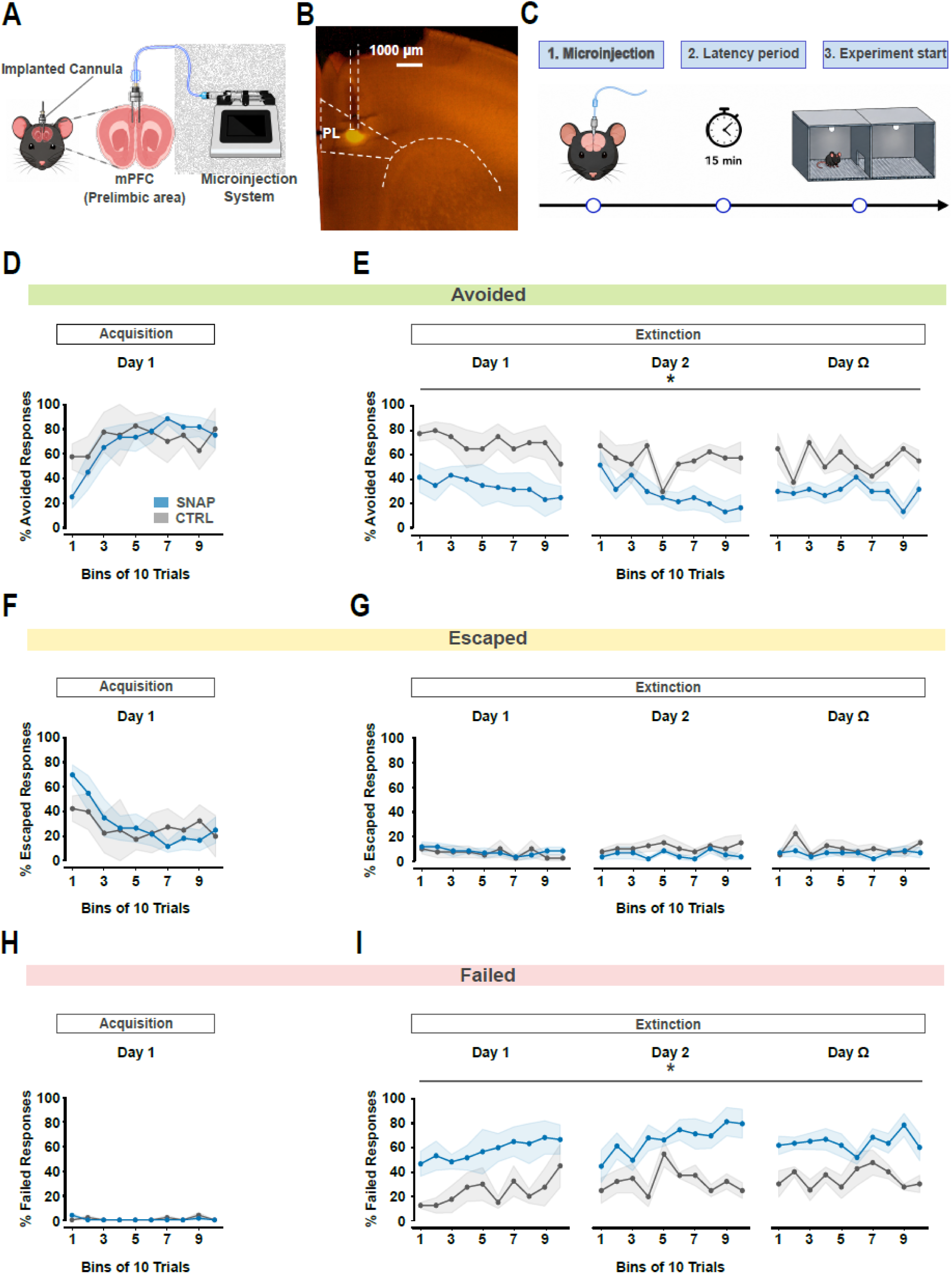
mPFC-targeted MCHR1 antagonism facilitates extinction of active avoidance. **a** Schematic illustration of the surgical procedure used for intracranial drug delivery. Mice were implanted with custom-made guide cannulae targeting the prelimbic region of the medial prefrontal cortex (mPFC). Drug microinjections were administered through injection needles connected to a microinfusion pump via polyethylene tubing. **b** Histological verification of the location of the targeted medial prefrontal cortex (mPFC; prelimbic cortex, PL). **c.** Schematic overview of the experimental timeline. Following intracranial microinjection, mice remained in their home cage for a 15 min incubation period to allow drug action before being placed into the behavioral apparatus to begin the task. **d** % Avoided responses across acquisition. Two-way repeated-measures ANOVA revealed a significant main effect of 10-trial bins (F(9, 72) = 5.16, p < 0.0001), with no significant main effect of Group factor (F(1,8) = 0.038, p = 0.85) and no significant interaction between factors (F(9, 72) = 1.80, p = 0.08). **e** % Avoided responses across extinction phase. Two-way repeated-measures ANOVA revealed a significant main effect of Group factor (F(1,8) = 9.55, p = 0.01), with significant main effect of 10-trial bins (F(29,232) = 2.19, p = 0.0008) and no significant interaction between factors (F(29, 232) = 1.25, p = 0.19). **f** % Escaped responses across acquisition. Two-way repeated-measures ANOVA revealed a significant main effect of 10-trial bins (F(9,72) = 4.652, p < 0.0001), with no significant main effect of Group factor (F(1,8) = 0.05, p = 0.82) and no significant interaction between factors (F(9,72) = 1.41, p = 0.20). **g.** % Escaped responses across extinction. Two-way repeated-measures ANOVA revealed no significant main effects of 10-trial bins (F(29, 232) = 0.85, p = 0.69) or Group factor (F(1,8) = 4.39, p = 0.07), and no significant interaction between factors (F(29, 232) = 0.79, p = 0.77). **h**. % Failed responses across acquisition. Two-way repeated-measures ANOVA revealed a significant main effects of 10-trial bins (F(9, 72) = 2.537, p = 0.014), no significant Group factor (F(1,8) = 0.29, p = 0.60), or interaction between factors (F(2.448, 19.59) = 2.22, p = 0.13). **i.** % Failed responses across extinction. Two-way repeated-measures ANOVA revealed a significant main effect of Group factor (F(1,8) = 10.45, p = 0.01), with significant main effect of 10-trial bins (F(29, 232) = 1.89, p = 0.0053) and no significant interaction between factors (F(29, 232) = 1.03, p = 0.42). Data are shown as mean ± s.e.m; n = 4 mice in control group (2 males and 2 females), n = 6 mice in SNAP group (4 males and 2 females).

The escaped and failed response analyses further specify the behavioral shift. Escaped responses changed during acquisition without a group effect (Fig. 3f; 10-trial bins, F(9,72) = 4.65, p < 0.0001; Group, F(1,8) = 0.05, p = 0.83), and during extinction the group effect did not reach significance (Fig. 3g; Group, F(1,8) = 4.39, p = 0.07). By contrast, failed responses were not different during acquisition (Fig. 3h; Group, F(1,8) = 0.29, p = 0.60), but were significantly increased by mPFC-targeted SNAP during extinction (Fig. 3i; Group, F(1,8) = 10.45, p = 0.01). Because shock was omitted during extinction, this increase in failed responses indicates reduced expression of the previously reinforced avoidance action, not increased exposure to shock. Together, these data show that local MCHR1 antagonism in the medial prefrontal region facilitates extinction by shifting behavior away from cue-driven avoidance when the cue no longer has aversive consequence.

### Systemic MCHR1 antagonism enhances mPFC responses during cue-guided avoidance

We next asked whether systemic MCHR1 antagonism changes excitatory mPFC activity during active avoidance. We recorded calcium activity from CaMKII-positive neurons in the prelimbic region of the mPFC and analyzed responses separately for avoided, escaped and failed trials (Fig. 4a,b). This outcome-specific analysis was important because mPFC activity during active avoidance can reflect threat-cue processing, action selection and movement-related components. A global increase in mPFC activity would therefore be less informative than a selective change during trials in which mice successfully used the cue to guide avoidance.

**Fig. 4.**
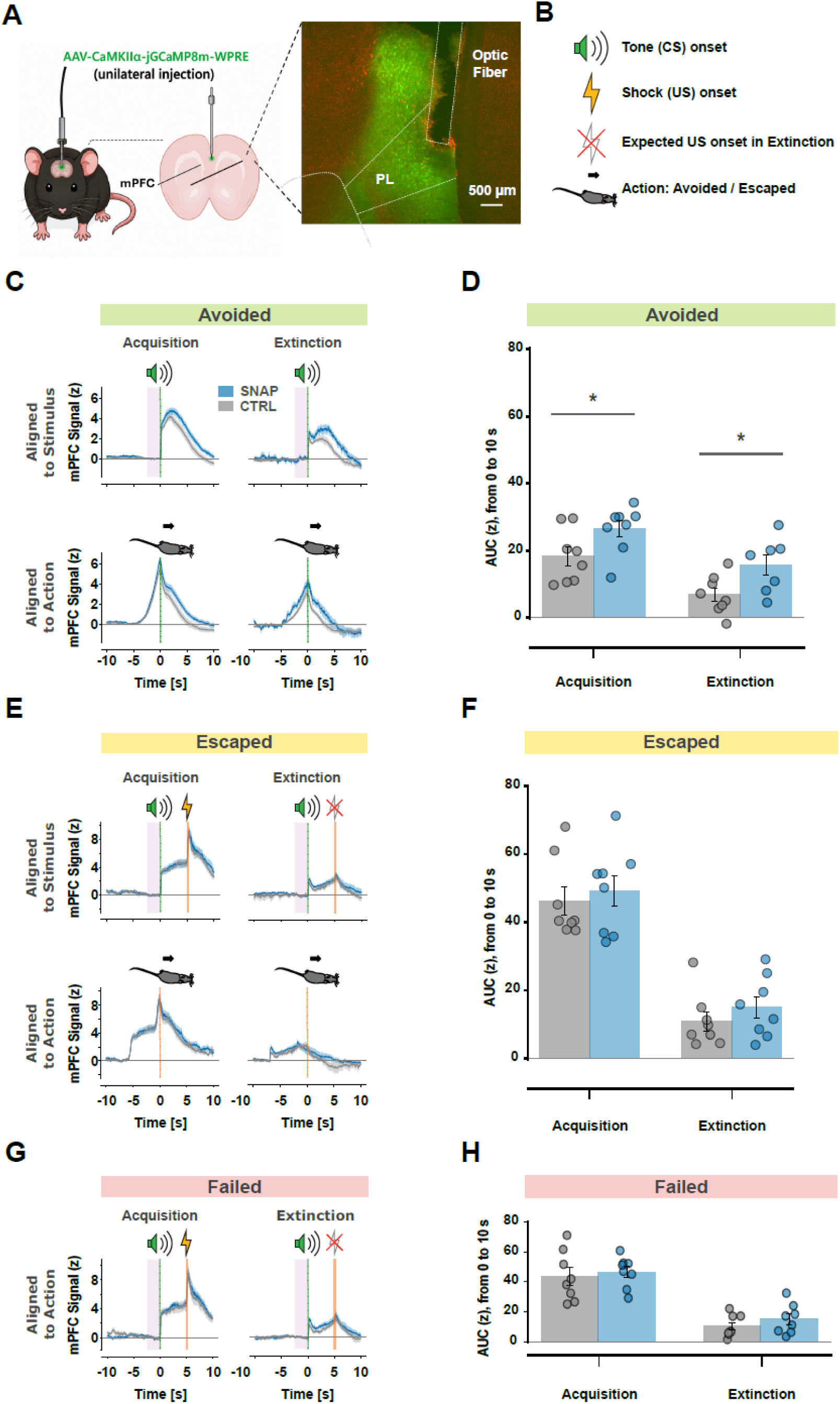
MCHR1 antagonism selectively alters excitatory mPFC activity during avoided trials. **a.** Schematic illustration of the surgical procedure for fiber photometry recordings. Mice received unilateral injections of the CamKIIα-jGCaMP8m-WPRE viral construct targeting the prelimbic region of the medial prefrontal cortex (mPFC), followed by unilateral optical fiber implantation for calcium signal recordings. The histological section verifies viral expression and correct fiber placement. **b.** Trial-event schematic used to align calcium signals to tone onset, shock onset or expected shock onset, and avoidance/escape actions. **c.** Avoided-trial photometry traces aligned to tone onset during acquisition (top left) and extinction (top right), and aligned to action onset during acquisition (bottom left) and extinction (bottom right). Traces are averaged across subjects. **d.** Area under the curve (AUC) for the 10 s window from tone onset for the avoided-trial traces shown in panel c. A significant difference between groups was found during acquisition (unpaired two-tailed Welch’s t-test, t = −2.18, p = 0.047) and extinction (unpaired two-tailed Welch’s t-test, t = −2.41, p = 0.04). **e.** Escaped-trial photometry traces aligned to tone onset during acquisition (top left) and extinction (top right), and aligned to action onset during acquisition (bottom left) and extinction (bottom right). Traces are averaged across subjects. **f.** AUC for the 10 s window from tone onset for the escaped-trial traces shown in panel e. No significant difference between groups was found during acquisition (unpaired two-tailed Welch’s t-test, t = −0.48, p = 0.64) or extinction (unpaired two-tailed Welch’s t-test, t = −0.98, p = 0.34). **g.** Failed-trial photometry traces aligned to tone onset during acquisition (top left) and extinction (top right). Traces are averaged across subjects. **h.** AUC for the 10 s window from tone onset for the failed-trial traces shown in panel g. No significant difference between groups was found during acquisition (unpaired two-tailed Welch’s t-test, t = −0.44, p = 0.67) or extinction (unpaired two-tailed Welch’s t-test, t = −1.09, p = 0.30). Data are shown as mean ± s.e.m; n = 8 mice in control group (5 males and 3 females), n = 8 mice in SNAP group (4 males and 4 females).

SNAP increased mPFC activity selectively during avoided trials. During acquisition, SNAP-treated mice showed significantly larger tone-onset responses than controls (Fig. 4c,d; Welch’s t-test, t = −2.18, p = 0.047). During extinction, SNAP also increased tone-onset activity during avoided trials (Fig. 4c,d; Welch’s t-test, t = −2.41, p = 0.04). Thus, although systemic MCHR1 antagonism did not impair acquisition behavior, it altered excitatory mPFC recruitment during successful cue-guided avoidance in both acquisition and extinction.

In contrast, SNAP did not significantly alter mPFC activity during escaped or failed trials. Escaped trials showed no group differences during acquisition or extinction (Fig. 4e,f; acquisition: t = −0.48, p = 0.64; extinction: t = −0.98, p = 0.34), and failed trials were similarly unchanged (Fig. 4g,h; acquisition: t = −0.44, p = 0.67; extinction: t = −1.09, p = 0.30). These results indicate that MCHR1 antagonism did not globally elevate mPFC activity across all behavioral outcomes. Rather, SNAP selectively enhanced excitatory mPFC activity during successful avoidance, consistent with increased prefrontal engagement when the warning cue is used to guide action.

### MCHR1 antagonism alters trial-history-dependent mPFC activity during extinction avoidance

We next asked whether the SNAP-enhanced mPFC response during avoidance depended on recent behavioral experience. Extinction is shaped not only by the current cue, but also by recent outcomes: whether avoidance was previously expressed, whether escape occurred, or whether non-avoidance was experienced without shock. This analysis was motivated by recent work showing that ACC/mPFC activity carries trial-history information, including previous choices and outcomes, separable from ongoing posture and movement (45). We therefore sorted CS-aligned mPFC activity by current trial outcome and by the immediately preceding trial outcome, yielding nine possible outcome transitions per phase (acquisition: Fig. 5a,c,e; extinction: Fig. 5g,i,k).

**Fig. 5.**
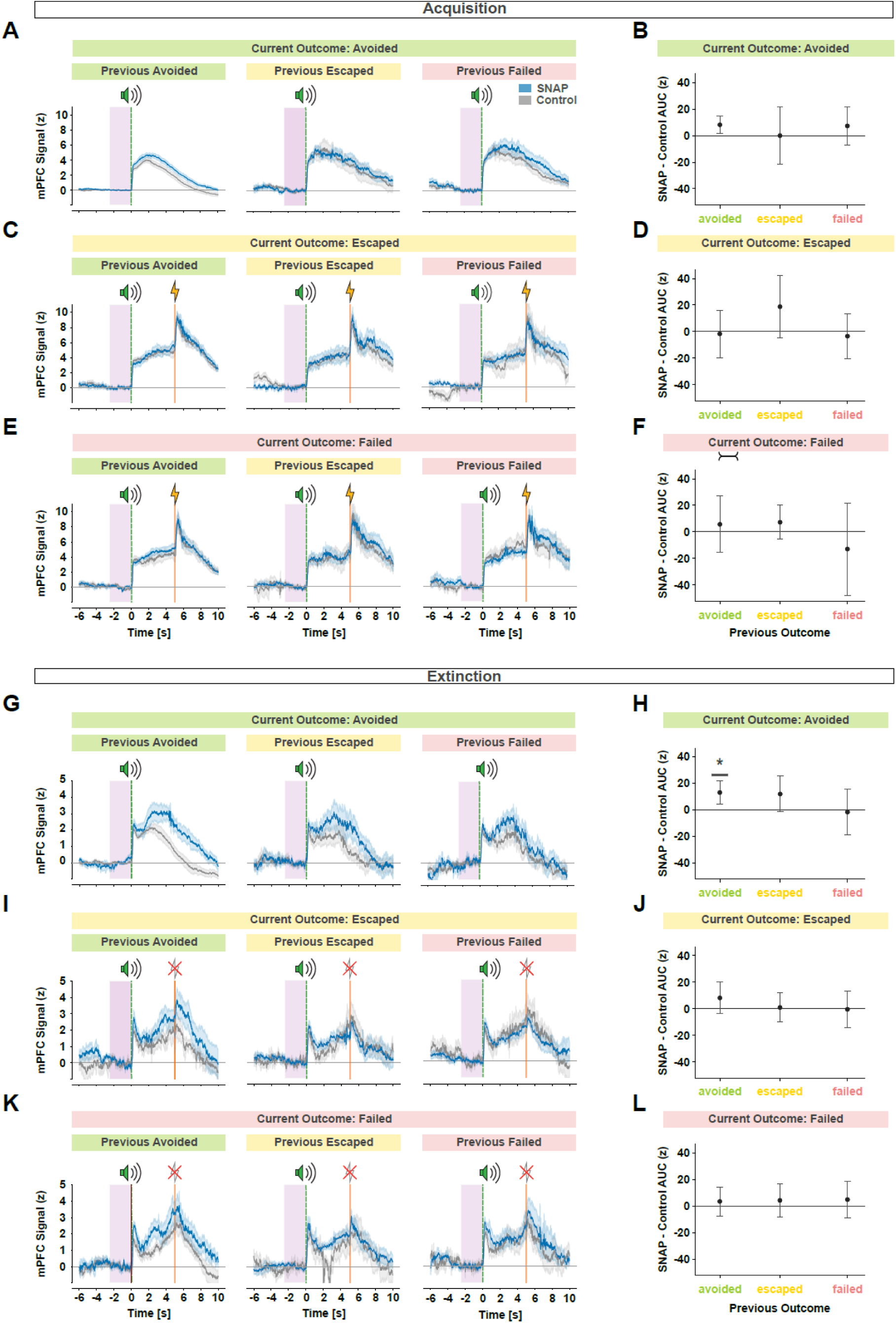
MCHR1 antagonism alters trial-history-dependent mPFC activity during extinction avoidance. **a, c, e, g, i, k** Trial-history-dependent photometry traces in Control and SNAP mice, aligned to tone onset and separated by phase, current outcome, and previous trial outcome. Traces are grouped according to the transition from the immediately preceding trial outcome to the current trial outcome. Acquisition trials are shown for current avoided (**a**), escaped (**c**), and failed (**e**) outcomes. Extinction trials are shown for current avoided (**g**), escaped (**i**), and failed (**k**) outcomes. Within each panel, traces are separated by previous avoided, previous escaped, or previous failed trials. Lines show subject-averaged group mean photometry traces for each outcome transition; shaded regions indicate SEM across mice. **b, d, f, h, j, l,** Model-based SNAP–Control contrasts from weighted trial-level AUC models fitted separately for each phase and current outcome. AUC was calculated over the 0–10 s window after tone onset. Models included Group, Previous outcome, categorical Day, and their interactions, with subject-balanced trial weights. Contrasts show the estimated SNAP–Control AUC difference for each previous-outcome condition, averaged across days. Error bars indicate 95% confidence intervals. P-values were obtained using CR2 cluster-robust small-sample correction with subject as the clustering variable. In extinction avoided trials (h), the SNAP–Control contrast averaged across extinction days was significant after previous avoided trials (estimate = 13.07 AUC, CR2-adjusted p = 0.03), but not after previous escaped trials (estimate = 11.96 AUC, CR2-adjusted p = 0.11) or previous failed trials (estimate = −1.66 AUC, CR2-adjusted p = 0.86).

Trial-level AUC values were analyzed using weighted linear models fitted separately by phase and current outcome, as described in Materials and Methods. During acquisition, failed trials showed a significant main effect of previous outcome (F(2, 4.66) = 10.86, p = 0.02), indicating that mPFC activity on failed acquisition trials depended on the preceding trial outcome. This effect was not accompanied by significant Group interactions, suggesting trial-history dependence shared across Control and SNAP mice rather than a SNAP-specific alteration.

During extinction, the strongest SNAP-dependent effect was observed for current avoided trials. In this condition, there was a significant Group × Previous outcome × Day interaction (F(4, 6.72) = 4.87, p = 0.04), indicating that SNAP altered the relationship between previous trial outcome and mPFC activity during extinction avoidance across days. Follow-up CR2-adjusted contrasts showed that the group difference reached significance only when the previous trial was also avoided. Averaged across extinction days, SNAP mice had higher AUC than Controls after previous avoided trials (estimate = 13.07 AUC, CR2-adjusted p = 0.03), but not after previous escaped trials (estimate = 11.96 AUC, CR2-adjusted p = 0.11) or previous failed trials (estimate = −1.66 AUC, CR2-adjusted p = 0.86).

Together, these results indicate that MCHR1 antagonism did not globally amplify mPFC activity across all trial histories. Instead, SNAP altered how recent avoidance history shaped excitatory mPFC activity during extinction, most clearly when animals expressed avoidance after a previous avoidance trial.

## Discussion

Adaptive avoidance is useful only as long as the environment still demands it. Once a warning cue no longer predicts harm, the same response that was previously protective must be suppressed. The present study identifies MCHR1 signaling as a regulator of this transition from persistent avoidance to extinction. Across systemic and mPFC-targeted pharmacological experiments, SNAP-94847 spared acquisition of active avoidance but facilitated extinction when tone presentations were no longer followed by shock. In adult mice, systemic MCHR1 antagonism reduced persistent avoidance during extinction and shifted behavior toward non-avoidance/failed responses, which in this phase reflects suppression of an outdated action rather than impaired defense. Local mPFC-targeted SNAP produced a parallel extinction phenotype, indicating that prefrontal MCHR1 signaling contributes to the persistence of cue-driven avoidance after the cue–shock contingency has changed. Thus, the central behavioral effect of MCHR1 antagonism is not a loss of avoidance capacity, but an acceleration of behavioral updating when avoidance is no longer required.

These findings fit with a modern view of active avoidance as an adaptive survival behavior that becomes maladaptive when it persists after danger has passed. Active avoidance is not equivalent to freezing or Pavlovian fear expression; it requires animals to use a threat-predictive cue to select an action that prevents harm, and then to suppress that action when the contingency changes (46) (6). Work in rodents and humans has shown that active avoidance can attenuate Pavlovian defensive responding and depends on interactions between prefrontal, amygdala and striatal circuits (20, 47) (26). Within this framework, the present data add a molecular and neuromodulatory layer: blocking MCHR1 does not abolish the capacity to avoid, but reduces the dominance of avoidance after the cue–shock contingency has changed. Thus, MCHR1 signaling appears to bias the system toward persistence of a previously successful defensive action, whereas MCHR1 antagonism facilitates the transition from action persistence to extinction.

The mPFC photometry data provide the mechanistic bridge between the behavioral phenotype and prefrontal circuit function. Because both the local infusion and photometry experiments targeted the prelimbic region of the mPFC, the data should be interpreted as implicating prelimbic medial prefrontal circuitry rather than the mPFC as an undifferentiated structure. We recorded calcium activity using a CaMKIIα-driven indicator, a strategy that enriches expression in cortical excitatory pyramidal neurons rather than broadly sampling all neuronal classes (48–50). These neurons constitute the principal cortical output population through which prelimbic mPFC can influence downstream amygdala and striatal circuits involved in avoidance. This choice is directly aligned with recent population-imaging work showing that mPFC excitatory activity contains avoidance-specific patterns that predict tone-induced avoidance actions and are separable from spontaneous movements with similar kinematics (28). In our study, SNAP selectively enhanced mPFC activity during avoided trials, whereas escaped and failed trials were largely unchanged. This selectivity argues against a global arousal or movement explanation and instead suggests that MCHR1 antagonism modifies excitatory prefrontal recruitment during successful cue-guided avoidance.

The timing of this photometry effect helps reconcile the behavioral and neural data. During acquisition, SNAP enhanced tone-onset mPFC responses during avoided trials despite no detectable impairment of avoidance acquisition. Thus, the absence of an acquisition phenotype does not mean that the MCH–mPFC circuit is inactive during learning. Rather, MCHR1 antagonism altered excitatory prefrontal processing while the tone–shock contingency was still valid, a condition in which rapid avoidance remained the appropriate behavioral strategy. During extinction, the enhanced tone-onset response is particularly informative because, once shock is omitted, the critical computation occurs when the cue begins and the animal must determine whether it still demands action. This pattern suggests that MCHR1 blockade enhances prefrontal cue evaluation when the auditory warning signal must be reinterpreted under a new, non-reinforced contingency, rather than simply increasing movement-related activity. This interpretation is consistent with active-avoidance work showing that prefrontal activity contributes to avoidance action selection and cannot be reduced to motor output alone (6, 26, 28).

The trial-history analysis provides evidence that MCHR1 antagonism affects adaptive updating rather than only current-trial avoidance. Extinction is built from recent experience: after each tone presentation without shock, the animal gains evidence about whether the learned avoidance strategy still has behavioral value. Using subject-balanced weighted models with small-sample correction, we found that during extinction, mPFC activity on avoidance trials was shaped by the immediately preceding trial outcome in a group- and day-dependent manner. The clearest SNAP-associated increase occurred when an avoided trial followed a previous avoided trial. This is important because repeated avoidance during extinction carries a different meaning than repeated avoidance during acquisition: the action is still being expressed, but it no longer prevents shock. SNAP therefore appears to enhance prefrontal processing of the very behavioral sequence that must be re-evaluated during extinction. Rather than uniformly increasing mPFC activity across all transitions, MCHR1 antagonism altered prefrontal activity associated with repeated avoidance under the new, non-reinforced contingency. This places the MCH system within a prefrontal computation of recent action history, consistent with evidence that ACC/mPFC populations encode previous choices and outcomes as trial-history variables separable from ongoing posture and movement (45).

This finding extends prior work placing MCH signaling in affective state, stress regulation, memory persistence and behavioral flexibility. MCHR1 antagonists, including SNAP-94847, have been studied in relation to anxiety-like and depressive-like behavior, stress responses and motivated behavior, indicating that the MCH system can influence more than metabolic state or sleep (40, 51–59). More recent circuit-level work showed that hypothalamic MCH neurons support fear extinction and long-term flexibility of fear behavior, while REM-active MCH neurons contribute to forgetting of hippocampus-dependent memories (38, 39). The present work moves this principle into instrumental defensive action by showing that MCHR1 antagonism facilitates extinction of active avoidance and links this behavioral effect to excitatory mPFC activity shaped by recent avoidance history. Thus, rather than merely reducing avoidance, MCHR1 antagonism appears to alter prefrontal engagement when recent experience indicates that an old avoidance rule has lost predictive value.

The aged cohort refines the boundary conditions of this mechanism. SNAP facilitated extinction on the first extinction day in aged mice, but this effect was not sustained across the full extinction protocol. Thus, MCHR1 antagonism can still initiate extinction-related behavioral change in aged animals, but sustained updating may require a circuit state that is less available with aging. This interpretation is consistent with broader evidence that aging alters cortical and hippocampal circuits supporting cognition, behavioral flexibility and adaptive updating (43, 44). One plausible explanation is that aging constrains the behavioral impact of MCHR1 blockade by altering mPFC excitability. Aging has been shown to impair extinction while redistributing mPFC excitability, including reduced excitability in extinction-promoting infralimbic neurons and increased excitability in prelimbic neurons that can oppose extinction (41). In such a circuit state, SNAP may transiently reduce expression of the learned avoidance strategy, but may be insufficient to sustain or consolidate extinction across repeated sessions. This interpretation remains a hypothesis because mPFC excitability and MCHR1 signaling were not directly measured in aged mice in the present study.

Several points define the scope of the conclusion. SNAP was administered during acquisition and extinction, so the study identifies how MCHR1 antagonism shapes the development and updating of avoidance rather than testing reversal of a fully established pathological avoidance state. This design is appropriate for the central question: whether the MCH system regulates adaptive behavior as cue-action-outcome contingencies are learned and revised. Future extinction-only manipulations will determine whether MCHR1 antagonism can also reverse stabilized avoidance after learning is complete. In parallel, projection-specific recordings or MCHR1 manipulations will be needed to determine whether the relevant mPFC signal is carried preferentially through mPFC-striatal, mPFC-amygdala, or other prefrontal output pathways. It will also be important to establish whether the same mechanism acts through direct MCHR1 signaling in prefrontal excitatory neurons or through local inhibitory and neuromodulatory intermediates that shape prefrontal population activity.

Together, these findings support a model in which MCHR1 signaling constrains prefrontal updating of instrumental avoidance. Such constraint may be adaptive in recently threatening or uncertain environments, where prematurely abandoning a protective action could be costly. Blocking MCHR1 increases excitatory mPFC engagement during successful cue-guided avoidance and alters extinction-related prefrontal activity according to recent avoidance history. In this model, MCHR1 antagonism facilitates extinction by reducing the persistence of a previously protective action while enhancing prefrontal processing when that action must be re-evaluated under a non-reinforced contingency. This positions MCH receptor signaling as a neuromodulatory mechanism capable of tuning the balance between defensive persistence and behavioral flexibility, linking survival-state signaling to mPFC-dependent control of adaptive action.

## Acknowledgments

This research was funded by ETH Grants (ETH-24 20-2 to D.P.-R.).

## Author contributions

D.P.R. conceived the study and designed the protocol with contributions from K.P.; Experimental setup K.P. and E.B.; K.P. performed research and L.O. assisted with experiments for Fig. 3; K.P. and D.P.R. analyzed data; Visualization K.P. and E.B.; D.P.R. wrote the manuscript with input from K.P., E.B. and D.B.

## Materials and Methods

### Experimental subjects

All animal procedures were conducted in accordance with Swiss Federal Food Safety and Veterinary Office regulations and were approved by the Cantonal Veterinary Office of Zürich (Animal Welfare Ordinance, SR 455.1). Adult (2–6 months old) and aged (17–21 months old) male and female C57BL/6 mice were used. Sex and sample size for each experiment are reported in the corresponding figure legends and Extended Data Table 1. Animals were housed under a reversed 12 h light/dark cycle in a temperature- and humidity-controlled environment (22 °C, 55% relative humidity) with ad libitum access to food and water. Experiments were performed during the dark phase of the cycle.

### Stereotaxic surgeries

Mice were anesthetized with isoflurane and placed in a stereotaxic frame (Kopf Instruments). Body temperature was maintained throughout surgery and animals received perioperative analgesia as described below. For all procedures, coordinates are given relative to bregma. In both intracranial pharmacology and fiber photometry experiments, the prelimbic region of the medial prefrontal cortex (PL/mPFC) was targeted.

For intracranial pharmacology experiments, mice were anesthetized with isoflurane (4–5% induction, 1–2% maintenance in oxygen) and received buprenorphine and local lidocaine at the incision site. Bilateral guide cannulae (RWD Life Science) were implanted above the PL/mPFC (AP, +1.94 mm; ML, ±0.3 mm; DV, −2.0 mm) and secured to the skull with dental cement. Animals were allowed to recover for at least 7 days before behavioral testing.

For fiber photometry experiments, mice were anesthetized with isoflurane and treated with carprofen (5 mg/kg, s.c.) for perioperative analgesia. A small craniotomy was made above the target site, and AAV1-CaMKIIα-jGCaMP8m-WPRE (titer ≥7 × 10^12 vg ml−1; Addgene) was injected unilaterally into the PL/mPFC using a Hamilton microsyringe. Virus (200 nL, undiluted) was delivered at 50 nL min−1 at AP, +1.8 mm; ML, +0.3 mm; DV, −2.0 mm. The injection needle was left in place for 5 min after infusion to minimize reflux. Fiber-optic implants (200 μm core diameter, 0.48 NA; Thorlabs) were implanted above the injection site (AP, +1.8 mm; ML, +0.3 mm; DV, −1.9 mm) and secured with dental cement. Animals were allowed to recover for at least 3 weeks before experiments.

### Drug preparation and administration

For systemic pharmacology, SNAP-94847 was dissolved in a vehicle containing 2% dimethyl sulfoxide (DMSO), (2-hydroxypropyl)-β-cyclodextrin (Merck) and phosphate-buffered saline (PBS). SNAP-94847 was administered intraperitoneally at 20 mg/kg, with injection volumes adjusted to body weight. Mice received injections 45 min before behavioral testing and remained in their home cages during the waiting period.

For intracranial microinjections, SNAP-94847 was dissolved in artificial cerebrospinal fluid (aCSF) containing 2% DMSO to a final concentration of 3 mg/ml. Mice were handled and habituated to the injection procedure before behavioral testing. Internal injectors were inserted into the implanted guide cannulae and connected via PE20 tubing to a Hamilton microsyringe. Infusions were delivered with a microinfusion pump (CMA 402) at 1 μL/min, with a total volume of 1 μL per hemisphere. Injectors were left in place for 4 min after infusion to allow diffusion and minimize backflow. Behavioral testing began after an additional 15 min incubation period in the home cage.

### Two-way active avoidance task

Two-way active avoidance was assessed in a shuttle box consisting of two identical compartments separated by a central doorway. The apparatus contained a stainless-steel grid floor connected to a shock generator (Coulbourn Instruments) and a custom auditory stimulus generator. The conditioned stimulus (CS) was an 85 dB tone, and the unconditioned stimulus (US) was a 0.3 mA foot shock.

The task consisted of acquisition and extinction phases. During acquisition, each trial began with a 5 s CS. If the animal did not shuttle to the opposite compartment during the CS, the CS was immediately followed by a 2 s US. A shuttle during the CS terminated the tone and prevented shock delivery; a shuttle during the 2 s US terminated the shock. Trials were classified according to the timing of the shuttle response: responses during the CS window were scored as avoided, responses during the US window were scored as escaped, and trials without a shuttle response during either window were scored as failed.

During extinction, the same temporal trial structure was used, but the US was omitted. Thus, avoided, escaped and failed responses were classified using the same CS and post-CS response windows defined during acquisition, allowing extinction behavior to be analyzed relative to the previously learned CS–US contingency.

Adult systemic and aged cohorts underwent 40-trial sessions during acquisition and extinction to maintain comparable behavioral conditions across age groups under identical systemic pharmacological manipulations; this session length was selected because aged mice did not reliably complete longer sessions. Cannulated mice underwent 100-trial sessions to minimize the number of repeated intracranial microinjections across training days and thereby reduce cumulative tissue damage associated with repeated infusions. Acquisition training continued until mice reached a predefined criterion of ≥70% avoidance responses across consecutive trial blocks. Extinction was then performed under no-shock conditions.

Behavioral performance was quantified as the percentage of avoided, escaped and failed responses across successive blocks of 10 trials.

### Fiber photometry recordings and analysis

Fiber photometry recordings were performed using a Doric Lenses system operating in lock-in mode with simultaneous excitation at 405 nm and 465 nm. The 465 nm channel was used to record calcium-dependent jGCaMP8m fluorescence, and the 405 nm channel was used as an isosbestic control for motion artifacts and calcium-independent signal fluctuations.

For signal processing, the 405 nm signal was linearly fitted to the 465 nm signal, and normalized fluorescence changes were calculated as %ΔF/F = ((F_465_ − F_405_fit) / F_405_fit) × 100, where F_465_ is the calcium-dependent fluorescence signal and F_405_fit is the fitted isosbestic control signal. Signal processing was adapted from previously published methods (Lerner et al., 2015; Gunaydin et al., 2014).

Photometry traces were aligned to task events according to trial outcome. Avoided trials were aligned to tone onset and tone offset. Escaped trials were aligned to tone onset, shock onset and shock offset during acquisition, and to the corresponding time windows during extinction. Failed trials were aligned to tone onset and to the shock-onset-equivalent time point. Area under the curve (AUC) was calculated over the 0–10 s window after each alignment event unless otherwise stated.

For trial-history analyses, trials were sorted according to the immediately preceding trial outcome and the current trial outcome.

Current avoided, escaped and failed trials were therefore analyzed as transition-defined events, including failed→avoided, escaped→avoided, avoided→avoided, avoided→escaped, escaped→escaped, failed→escaped, avoided→failed, escaped→failed and failed→failedtransitions. Acquisition and extinction phases were analyzed separately.

### Histology and verification of cannula and fiber placement

After completion of experiments, animals were deeply anesthetized with pentobarbital and transcardially perfused with PBS (pH 7.4), followed by 4% paraformaldehyde (PFA) in PBS. Brains were removed, post-fixed overnight in 4% PFA at 4 °C and cryoprotected in 30% sucrose in PBS overnight. Tissue was frozen on dry ice and sectioned coronally at 50 μm using a cryostat. Sections were imaged with a fluorescence microscope (Eclipse Ti2, Nikon) to verify cannula and fiber placement.

Cannula targeting and injector patency were verified using fluorescein isothiocyanate–dextran (FITC–dextran; CAS no. 60842-46-8; Sigma-Aldrich) infused through the same injector configuration. FITC–dextran fluorescence confirmed delivery to the targeted prelimbic mPFC region and the expected local infusion field around the injector tip. Animals with cannula or fiber placements outside the intended target region were excluded from analysis.

### Data analysis and statistics

Statistical analyses and graphical visualizations were performed and generated using GraphPad Prism 11 and Python 3.9. Data are presented as mean ± s.e.m. unless otherwise stated. Exact sample sizes, experimental units and statistical tests are reported in the figure legends. Statistical significance was set at P < 0.05.

Behavioral data were analyzed using two-way repeated-measures ANOVA or mixed-effects ANOVA, as appropriate for repeated measurements across trial bins across days. In case of violations Greenhouse-Geisser correction was used. Post hoc comparisons and multiple-comparison corrections, when applied, are specified in the corresponding figure legends. Area under the curve analyses of outcome-specific photometry data in Fig. 4 were performed using unpaired two-tailed Welch’s t-tests. Trial-history analyses in Fig. 5 were analyzed using weighted trial-level linear models fitted separately by phase and current outcome. Models included Group, Previous outcome, categorical Day and their interactions. For each phase and current outcome, trial-level AUC values were modeled as AUC ∼ Group × Previous outcome × Day, with Day treated as a categorical factor. To prevent subjects or trial types with more traces from dominating the analysis, each trial was weighted by the inverse number of trials within each Subject × Phase × Day × Current outcome × Previous outcome cell, so that each cell contributed a total weight of 1. Statistical inference used CR2 cluster-robust small-sample correction with subject as the clustering variable. Significant omnibus effects were followed by model-based SNAP-Control contrasts within previous-outcome conditions.

Male and female mice were initially analyzed separately to assess potential sex-dependent effects. Because no significant sex differences were detected in the analyzed behavioral datasets, data from both sexes were pooled within experimental groups for the final analyses. P values are reported exactly where possible. Significance indicators are defined as *P < 0.05, **P < 0.01, ***P < 0.001 and ****P < 0.0001; NS, P > 0.05.

